# Meclizine rescues cardiac function and mitochondrial ultrastructure by ATP- and glycolysis-independent mechanisms in a genetic model of mitochondrial energy dysfunction

**DOI:** 10.1101/2025.09.03.674000

**Authors:** Nasab Ghazal, Banjamin Huang, Luke J. Shoemaker, Victor Faundez, Jennifer Q. Kwong

**Affiliations:** Graduate Program in Biochemistry, Cell and Developmental Biology; Graduate Division of Biological and Biomedical Sciences, Emory University, Atlanta, GA, USA; Division of Pediatric Cardiology, Department of Pediatrics, Emory University School of Medicine, and Children’s Healthcare of Atlanta, Atlanta, GA, USA; Department of Cell Biology, Emory University School of Medicine, Atlanta, GA, USA

## Abstract

Primary mitochondrial cardiomyopathies are an unmet clinical challenge, as there are no therapies that directly address the underlying mitochondrial dysfunction. We previously reported that the cardiomyocyte-specific deletion of the mitochondrial phosphate carrier (SLC25A3), which imports phosphate required for ATP synthesis, produces a model of mitochondrial cardiomyopathy in which total cardiac ATP levels are preserved despite defective mitochondrial ATP production. This was accompanied by increased glycolytic activity and reduced mitochondrial flux, leading us to hypothesize that pharmacologically enhancing glycolysis might be protective when the mitochondrial energy machinery is intrinsically impaired. To test this, we turned to meclizine, an FDA-approved antihistamine previously shown to shift metabolism toward glycolysis. Chronic meclizine treatment in SLC25A3-deficient mice attenuated cardiac hypertrophy, improved systolic function, and restored mitochondrial ultrastructure. Unexpectedly, meclizine suppressed glycolytic enzyme expression and reduced lactate accumulation, suggesting that meclizine does not induce a glycolytic shift in SLC25A3-deleted hearts. Instead, proteomic and functional analyses revealed preservation of mitochondrial cristae architecture via MICOS upregulation and improved NAD^+^/NADH homeostasis through uncoupled electron flux and NAD^+^ regeneration. Together, these findings identify meclizine as a clinically approved compound that promotes cardioprotection in mitochondrial disease not by driving glycolysis, but by preserving mitochondrial membrane organization and redox balance, highlighting mitochondrial quality and NAD^+^ redox homeostasis as therapeutic targets for primary mitochondrial cardiomyopathies.

## INTRODUCTION

Primary mitochondrial disorders are a group of individually rare conditions that collectively affect approximately 1 in 5,000 people, making them a common type of inherited metabolic disease (Ng & Turnbull, 2016). These heterogeneous conditions result from mutations in nuclear or mitochondrial DNA-encoded mitochondrial proteins that impair mitochondrial oxidative phosphorylation (OXPHOS) and energy production, often impact tissues with high energy demands (Anan *et al*, 1995; Meyers *et al*, 2013; Niyazov *et al*, 2016). In the heart, primary mitochondrial disorders can manifest as cardiomyopathies characterized by cardiac hypertrophy, contractile dysfunction, mitochondrial structural abnormalities, mitochondrial hyperproliferation, and impaired OXPHOS (Gorman *et al*, 2016; Holmgren *et al*, 2003; Limongelli *et al*, 2017; Marin-Garcia & Goldenthal, 2002; Rosca *et al*, 2013). Despite their clinical impact, to date there are no targeted therapies for mitochondrial cardiomyopathies, and current management relies on standard heart failure treatments that do not address the underlying mitochondrial dysfunction (Meyers *et al*., 2013). This therapeutic limitation underscores the need to define how the heart responds to mitochondrial energy dysfunction and to identify interventions that can preserve cardiac function and ATP homeostasis when the mitochondrial energy production machinery is compromised.

A central unanswered question is whether cellular energy homeostasis can be maintained despite intrinsic defects in OXPHOS and mitochondrial ATP synthesis. In prior studies, we developed a mouse model with tamoxifen-inducible cardiomyocyte-specific deletion of the mitochondrial phosphate carrier (SLC25A3; *Slc25a3^fl/flxMCM^* mice) (Kwong *et al*, 2014). This mitochondrial inner membrane transporter regulates mitochondrial import of inorganic phosphate, a requisite substrate for mitochondrial ATP synthesis (Kolbe *et al*, 1984; Palmieri, 2004). Deletion of SLC25A3 in the heart leads to impaired mitochondrial ATP synthesis and recapitulates clinical features of human phosphate carrier deficiency (MPCD, OMIM #610773), including cardiac hypertrophy, mitochondrial hyperproliferation, mitochondrial dysfunction, and contractile dysfunction (Bhoj *et al*, 2015; Mayr *et al*, 2007; Mayr *et al*, 2011). Surprisingly, despite mitochondrial ATP synthesis defects, SLC25A3-deleted hearts exhibit preserved total ATP levels (Kwong *et al*., 2014). We also noted increased cardiac glucose uptake, enhanced glycolytic enzyme expression, and upregulation of pyruvate dehydrogenase kinase 4 (PDK4) (Kwong *et al*., 2014). Those findings prompted us to postulate that an overall metabolic shift away from mitochondrial OXPHOS toward glycolytic ATP production may provide an adaptive and protective mechanism to maintain energy balance under conditions of intrinsic mitochondrial energetic stress.

In line with this, we considered that a pharmacologic approach to enhance glycolytic flux may yield cardioprotection against mitochondrial ATP synthesis defects. One intriguing drug candidate is meclizine, an FDA-approved antihistamine that can promote a metabolic shift toward glycolysis by inhibiting mitochondrial respiration and altering phospholipid metabolism (Gohil *et al*, 2010; Gohil *et al*, 2013; Houston *et al*, 2025). Indeed, meclizine confers protection in contexts of mitochondrial stress, including ischemia-reperfusion injury and neurodegeneration, by promoting glycolytic metabolism (Gohil *et al*, 2011; Gohil *et al*., 2010; Kishi *et al*, 2015). Thus, we hypothesized that meclizine might enhance glycolytic compensation and preserve cardiac function in primary mitochondrial cardiomyopathy due to defects in mitochondrial energy production like those due to SLC25A3 dysfunction.

To test this hypothesis, we deployed meclizine treatment in our established SLC25A3 cardiomyocyte-specific deletion model and assessed effects on cardiac and mitochondrial structure and function. Contrary to previous studies reporting that meclizine enhances glycolytic flux, we found no evidence of increased glycolysis in the hearts of meclizine-treated mice. Instead, our findings reveal an unexpected pathway by which long-term systemic meclizine treatment confers cardioprotection: by stabilizing mitochondrial structure and restoring NAD^+^ redox balance even when the mitochondrial ATP synthesis machinery is compromised. Together, these findings challenge the prevailing view of meclizine’s mechanism of action in vivo, and suggest that clinically approved metabolic modulators like meclizine may protect the energy-deprived heart in mitochondrial disease by enhancing mitochondrial health rather than directly restoring ATP synthesis.

## RESULTS

### Meclizine attenuates cardiac hypertrophy and enhances cardiac function in the SLC25A3 deletion model of mitochondrial cardiomyopathy

Given prior reports suggesting that meclizine can promote glycolysis (Gohil *et al*., 2011; Gohil *et al*., 2010; Gohil *et al*., 2013), we hypothesized that meclizine treatment might preserve cardiac function in the setting of mitochondrial energy dysfunction. To test this hypothesis, we examined the effects of meclizine in mice with tamoxifen-inducible cardiomyocyte-specific SLC25A3 deletion (*Slc25a3^fl/flxMCM^*mice), a well-established model of mitochondrial cardiomyopathy (Kwong *et al*., 2014). To induce SLC25A3 deletion, tamoxifen was administered intraperitoneally to 8-week-old *Slc25a3^fl/flxMCM^* mice; *Slc25a3^fl/fl^*littermates served as controls (Figure 1A). Concurrent with the initiation of tamoxifen administration, mice of both genotypes were randomized to receive either vehicle (10% Kolliphor) or meclizine (100 mg/kg/day) by oral gavage. Vehicle or meclizine treatments were continued daily for 12 weeks to evaluate the therapeutic potential of meclizine in the context of chronic mitochondrial energy deficiency.

**Figure 1.**
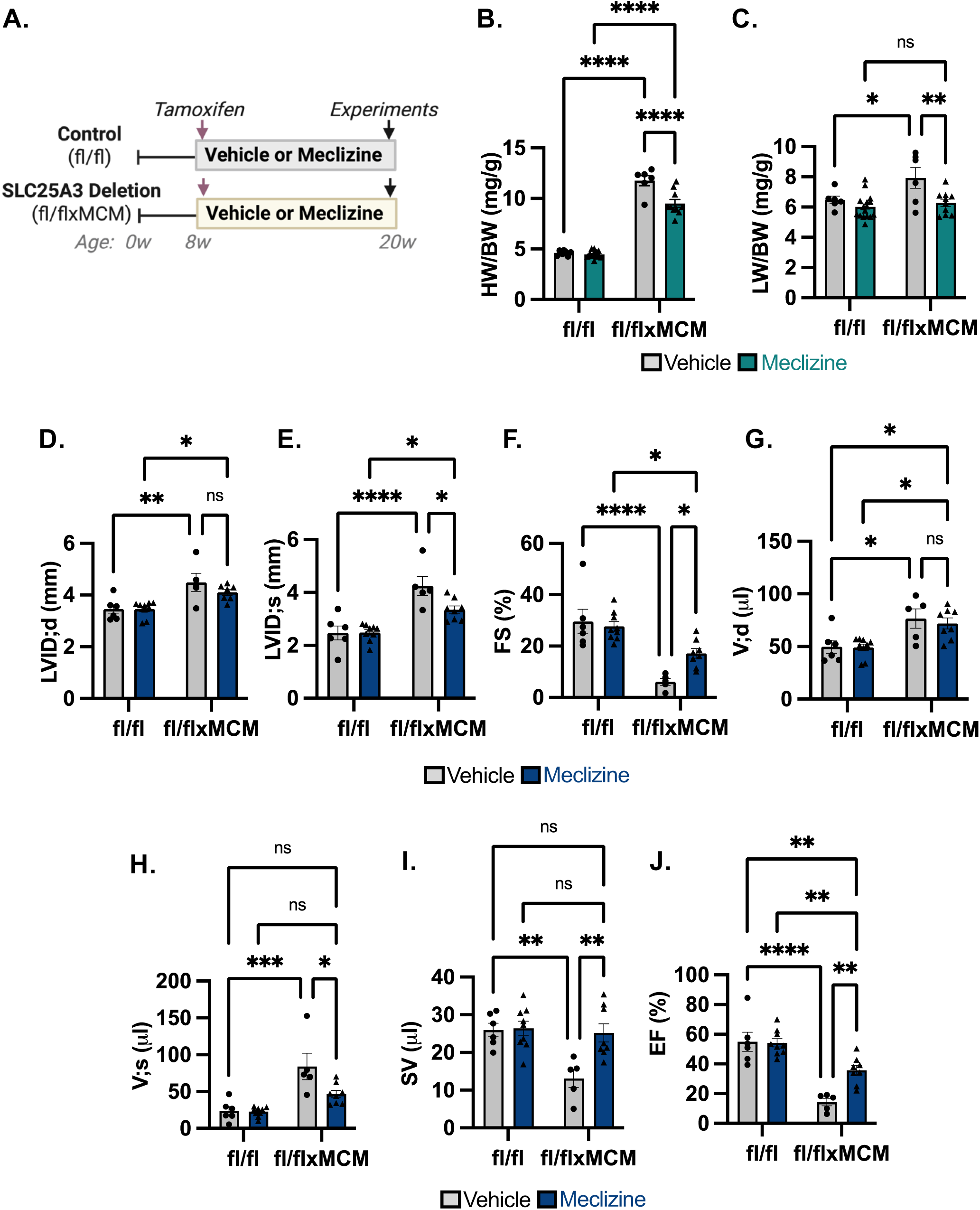
Systemic meclizine treatment attenuates pathological remodeling and preserves cardiac function in the SLC25A3 deletion model of mitochondrial cardiomyopathy. (A) Schematic of the treatment protocol. Cardiac-specific SLC25A3 knockout mice were generated using tamoxifen inducible Cre recombinase (25 mg/kg/day intraperitoneally for 5 days). Mice were treated with vehicle or meclizine (100 mg/kg/day by oral gavage) concurrent with the tamoxifen initiation of *Slc25a3* deletion, and hearts were collected at 12 weeks post deletion for various assays. (B) Heart weight-to-body weight ratio (HW/BW). (C) Lung weight-to-body weight ratio (LW/BW). (D-E) Echocardiographic measurements of left ventricular internal diameter in diastole (LVID; d) and systole (LVID; s). (F) Fractional shortening (FS), (G) end-diastolic volume (V; d), (H) end-systolic volume (V; s), (I) stroke volume (SV), and (J) ejection fraction (EF) was assessed 12 w post-tamoxifen/meclizine. Data are shown as mean ± SEM. Two-way ANOVA was used to determine statistical significance. \**p* < 0.05, \*\**p* < 0.01, \*\*\**p* < 0.001, \*\*\*\**p* < 0.0001. Diagram created with BioRender.

Consistent with our previous observations (Kwong *et al*., 2014), vehicle-treated *Slc25a3^fl/flxMCM^* mice developed pronounced cardiac hypertrophy and pulmonary edema, reflected by significantly elevated heart weight-to-body weight (HW/BW) and lung weight-to-body weight (LW/BW) ratios compared to vehicle-treated *Slc25a3^fl/fl^*controls (Figure 1B,C). In contrast, meclizine-treated *Slc25a3^fl/flxMCM^*mice exhibited a significant reduction in both HW/BW and LW/BW ratios relative to vehicle-treated *Slc25a3^fl/flxMCM^* mice, indicative of attenuation of pathological cardiac remodeling and hypertrophy (Figure 1B,C).

Echocardiographic analysis following completion of the meclizine and vehicle dosing regimens demonstrated that meclizine significantly improved systolic function in meclizine versus vehicle treated *Slc25a3^fl/flxMCM^*mice. Although left ventricular internal diameter at diastole (LVID;d) remained comparable between groups (Fig. 1D), meclizine-treated *Slc25a3^fl/flxMCM^* mice showed a significant reduction in left ventricular internal diameter at systole (LVID;s, Fig. 1E), accompanied by increased fractional shortening (%FS, Fig. 1F). End-systolic volume (V;s) was similarly reduced with meclizine treatment (Fig. 1H), while end-diastolic volume (V;d) was unchanged (Fig. 1G), resulting in restoration of stroke volume (Fig. 1I) and a significant increase in ejection fraction (Fig. 1J). Together, these data demonstrate that systemic meclizine administration reduces pathological cardiac remodeling and improves contractile performance in the setting of cardiomyocyte-specific mitochondrial energy deficiency due to SLC25A3 loss.

### Meclizine treatment does not alter the cardiac proteome in control mice

To define the mechanisms by which meclizine treatment conferred protection in the SLC25A3 deletion mitochondrial cardiomyopathy model, we performed tandem mass tag (TMT)-based quantitative proteomic profiling (Figure 2A) on cardiac tissue from four experimental groups: tamoxifen-treated *Slc25a3^fl/flxMCM^* and *Slc25a3^fl/fl^* control mice administered either meclizine or vehicle (Figure 1A). Across all samples, profiling identified 20530 peptides corresponding to 2901 proteins. After filtering for proteins with complete quantification across all samples and applying a false discovery rate (FDR) threshold of <1%, 2,803 proteins were retained for downstream analysis. We determined differentially expressed proteins (DEPs) using a Student’s t-test with Benjamini-Hochberg correction (log_2_ fold change > 0.58 and adjusted p-value < 0.05).

**Figure 2.**
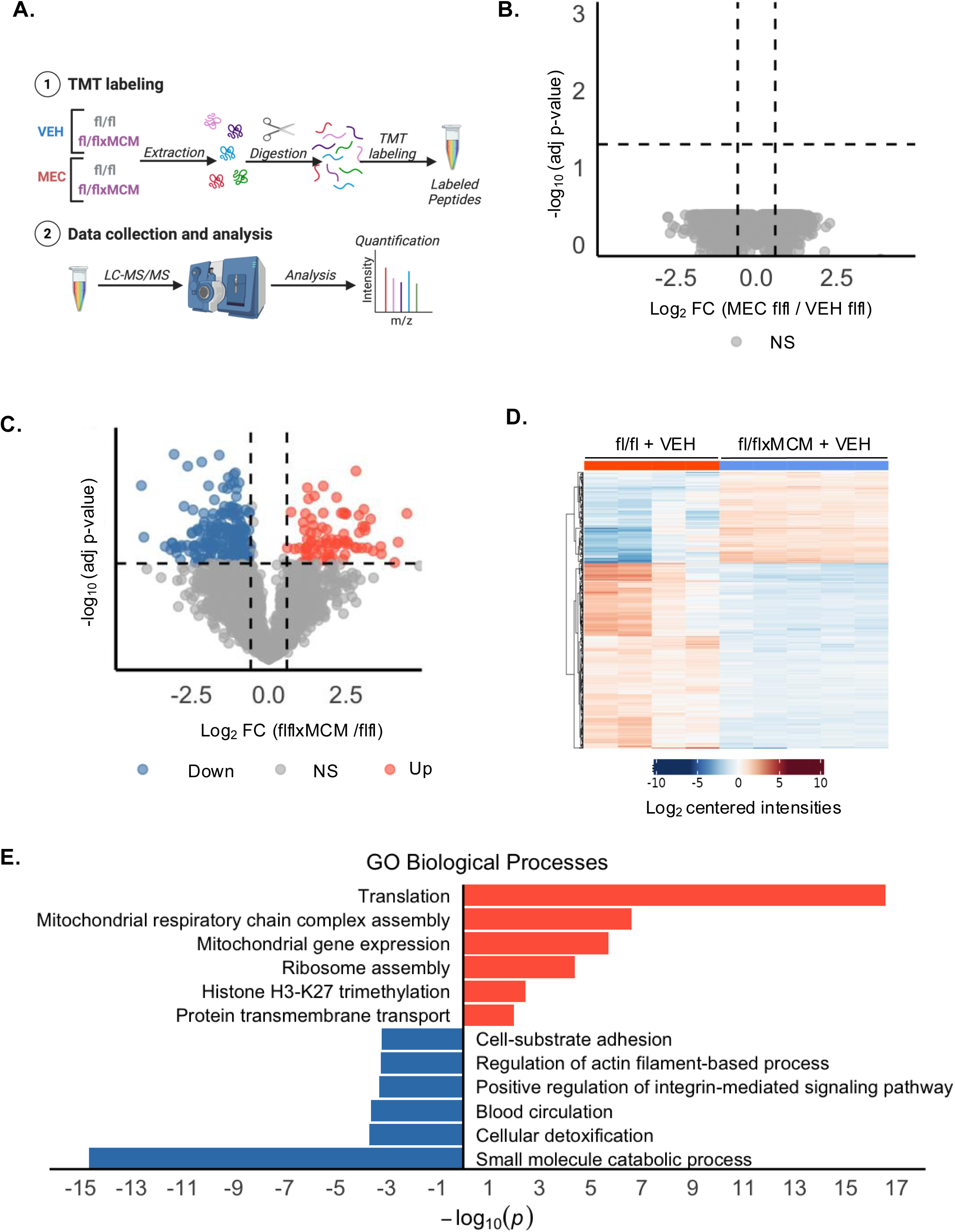
Proteomic profiling reveals coordinated translational and metabolic remodeling with loss of SLC25A3 in the heart. (A) Experimental design for TMT-based quantitative proteomic analysis of cardiac tissue from meclizine- or vehicle-treated *Slc25a3^fl/flxMCM^* (cardiac-specific knockout) and *Slc25a3^fl/fl^* (control) mice. (B) Volcano plot highlighting significantly differentially expressed proteins (DEPs) (MEC vs: VEH treated *Slc25a3^fl/fl^* controls) (log₂ fold change > 0.58 or < –0.58, adjusted p < 0.05) gray dots represents proteins that are not significantly altered (n=4/group). (C) Volcano plot highlighting significantly differentially expressed proteins (DEPs) (vehicle treated: *Slc25a3^fl/flxMCM^* vs. *Slc25a3^fl/fl^* hearts) (log₂ fold change > 0.58 or < –0.58, adjusted p < 0.05). Blue and red dots represent proteins that are significantly downregulated and upregulated, respectively (n=4 *Slc25a3^fl/fl^*, n=5 *Slc25a3^fl/flxMCM^*). (D) Heatmap of DEPs (vehicle treated: *Slc25a3^fl/flxMCM^* vs. *Slc25a3^fl/fl^*) where rows are individual proteins and columns are different replicates; red indicates increased expression while blue indicates reduced expression (Supplemental table 1). (E) ClueGo analysis of vehicle treated *Slc25a3^fl/flxMCM^* vs. vehicle treated *Slc25a3^fl/fl^* DEPs. The GO Biological Process was queried with a threshold Bonferroni-corrected *p* value <0.05 (Supplemental table 2). A subset of significant processes is represented by graphing -log_10_ (p); the red bars represent upregulated processes and blue represent downregulated processes. Significant DEPs were determined (log₂ fold change > 0.58 or < –0.58, adjusted p < 0.05) by moderated t-test with Benjamini-Hochberg correction for multiple testing. Diagram created with BioRender.

To specifically assess the effects of meclizine on the healthy heart, we compared proteomic profiles of vehicle- and meclizine-treated *Slc25a3^fl/fl^* control hearts (Figure 2B, Supplemental Table 1). No proteins met significance thresholds for differential expression, indicating that meclizine does not alter the cardiac proteome under non-pathological conditions.

### Proteomic profiling reveals critical changes to translation and ribosomal pathways in the SLC25A3 deleted heart

While the physiological consequences of SLC25A3 deficiency in the heart are characterized (Kwong *et al*., 2014), the cardiac pathways disrupted by loss of mitochondrial phosphate transport remain incompletely defined (Peoples *et al*, 2021). To address this, we compared the cardiac proteomes of vehicle-treated *Slc25a3^fl/flxMCM^* and *Slc25a3^fl/fl^* control mice. Comparative analysis revealed 226 proteins significantly (log2 fold change > 0.58 and adjusted p-value < 0.05) upregulated, and 83 proteins downregulated in the knockout hearts (Figure 2C and 2D, Supplemental Table 1). Gene ontology enrichment analysis of biological processes of proteins upregulated with SLC25A3 deletion revealed a significant overrepresentation of proteins annotated to pathways related to cytoplasmic translation (GO:0006412, p = 2.71E-17, group p value Bonferroni corrected) and mitochondrial respiratory chain complex assembly (GO:0033108, p = 2.6E-07, group p value Bonferroni corrected) and ribosome assembly (GO:0042255, p =4.37E-05, group p value Bonferroni corrected) (Figure 2E, Supplemental Table 2). These findings suggest activation of adaptive programs promoting mitochondrial protein synthesis and organelle proliferation in response to impaired energy production, consistent with the previously reported mitochondrial hyperproliferation in this model (Kwong *et al*., 2014).

Conversely, proteins involved in small molecule catabolic processes (GO:0044282, 2.16E-15, group p value Bonferroni corrected), including organic acid (GO:0016054, 2.16E-15, group p value Bonferroni corrected), amino acid (GO:0006520, 2.16E-15, group p value Bonferroni corrected), branched-chain amino acid (GO:0009083, 2.16E-15, group p value Bonferroni corrected), and fatty acid metabolism (GO:0006631, 2.16E-15, group p value Bonferroni corrected), as well as proteins involved with extracellular matrix and cell adhesion pathways (GO:0031589, 6.14E-04, group p value Bonferroni corrected) were significantly downregulated with SLC25A3 deletion (Figure 2E, Supplemental Table 2). This pattern may reflect a shift toward anabolic remodeling to support hypertrophic growth or mitochondrial biogenesis, or a broader disruption of catabolic flux driven by energy deficiency. Together, these findings suggest that loss of SLC25A3 triggers coordinated remodeling of translational, mitochondrial, and metabolic networks in the heart in response to impaired mitochondrial energy production.

### Meclizine induces mitochondrial remodeling independent of core SLC25A3 deletion-driven proteomic changes

To define the molecular targets of meclizine, we compared proteomic profiles of meclizine-treated *Slc25a3^fl/flxMCM^* hearts to meclizine-treated *Slc25a3^fl/fl^* controls. This analysis identified 991 differentially expressed proteins (DEPs), including 360 upregulated and 634 downregulated proteins (Figure 3A and 3B, Supplemental Table 1).

**Figure 3.**
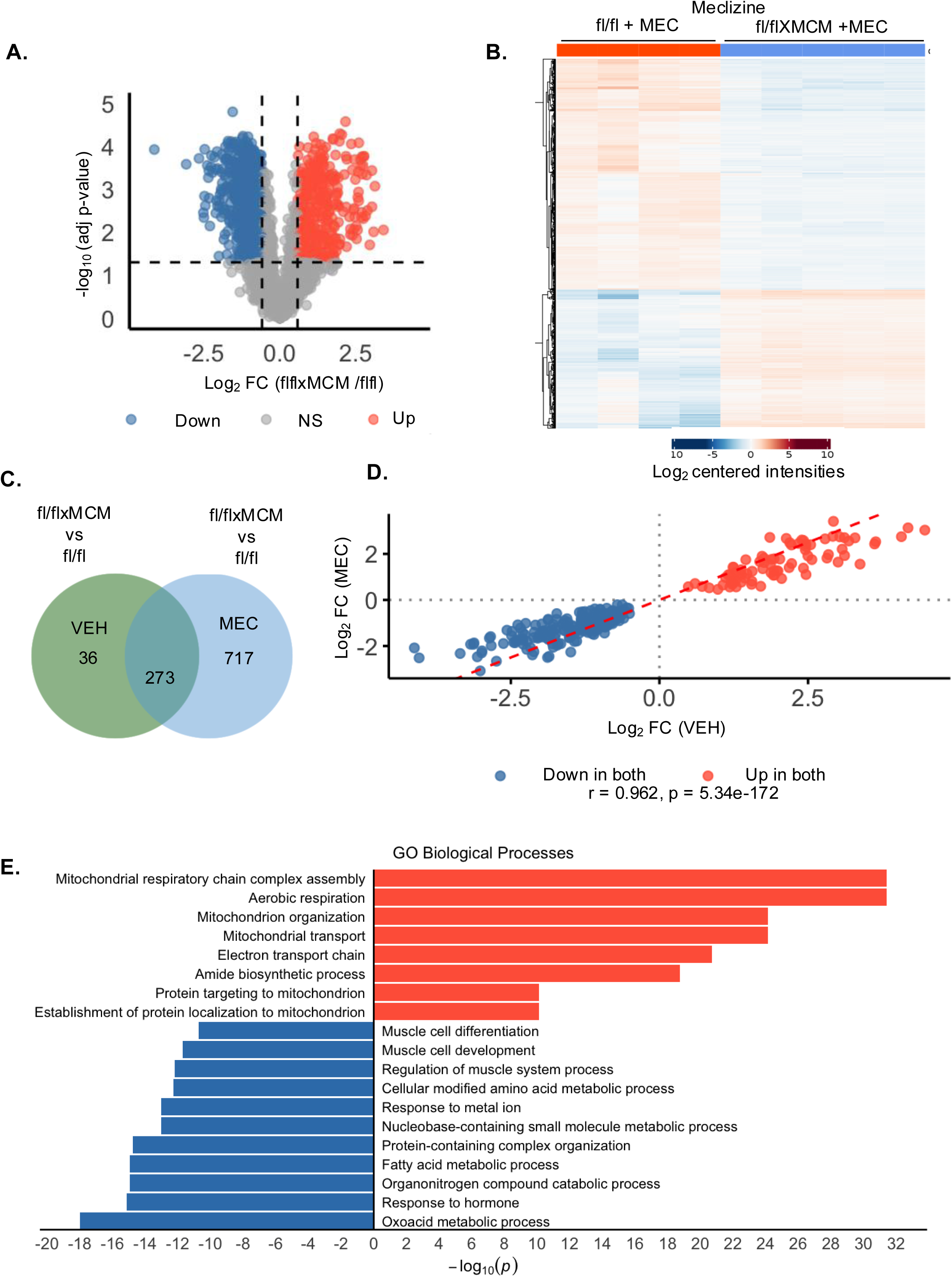
Meclizine induces a distinct proteomic program linked to mitochondrial remodeling and metabolic stress regulation. (A) Volcano plot of the DEPs (meclizine treated: *Slc25a3^fl/flxMCM^* vs. *Slc25a3^fl/fl^*) (log₂ fold change > 0.58 or < –0.58, adjusted p < 0.05). Blue and red dots represent proteins that are significantly downregulated and upregulated, respectively (n=4 *Slc25a3^fl/fl^*, n=5 *Slc25a3^fl/flxMCM^*). (B) Heatmap of DEPs of (meclizine treated: *Slc25a3^fl/flxMCM^* vs. *Slc25a3^fl/fl^*), where rows are individual proteins and columns are different replicates. Red indicates increased expression, while blue indicates reduced expression (Supplemental table 1). (C) Venn diagram showing the overlap of DEPs from vehicle and meclizine treated hearts. (D) Pearson’s correlation plot of log₂ fold changes for the 273 proteins altered in both meclizine-treated (*Slc25a3^fl/flxMCM^* vs. *Slc25a3^fl/fl^*) and vehicle-treated (*Slc25a3^fl/flxMCM^* vs. *Slc25a3^fl/fl^*) hearts. Each point represents a shared DEP, colored by direction of change (red = up in both, blue = down in both) (r = 0.962) (Supplemental table 3). (E) ClueGo analysis of DEPs in meclizine treated hearts (*Slc25a3^fl/flxMCM^* vs *Slc25a3^fl/fl^*). The GO Biological process database was queried with a threshold Bonferroni-corrected *p*-value< 0.05 (Supplemental table 4). Significant DEPs were determined (log₂ fold change > 0.58 or < –0.58, adjusted p < 0.05) by moderated t-test with Benjamini-Hochberg correction for multiple testing.

To determine whether meclizine alters the same proteomic pathways caused by SLC25A3 deletion or acts on a distinct set of targets, we compared differentially expressed proteins (DEPs) identified in meclizine-treated knockout hearts to those from vehicle-treated knockout hearts. Out of 991 DEPs in meclizine-treated hearts, 273 were also altered in vehicle-treated knockouts (Figure 3C). For these shared proteins, log₂ fold changes were highly correlated between groups (r = 0.962; p < 5.34e-172; Figure 3D, Supplemental Table 3), showing that meclizine does not broadly reverse the SLC25A3-associated proteomic signature. Instead, most changes with meclizine occur in proteins not altered by vehicle treatment, indicating that meclizine engages an additional, independent set of targets.

To gain insight into the functional consequences of these meclizine-specific proteomic changes, we performed GO enrichment analysis of biological processes on the 717 DEPs unique to meclizine-treated *Slc25a3^fl/flxMCM^*hearts (280 upregulated proteins, 437 downregulated proteins; Figure 3E, Supplemental Table 4). Upregulated categories were enriched for mitochondrial pathways, including respiratory chain complex assembly (GO:0033108, 6.96E-25, group p value Bonferroni corrected), aerobic respiration (GO:0009060, 3.77E-32, group p value Bonferroni corrected), and mitochondrial organization (GO:0007005, 6.96E-25, group p value Bonferroni corrected). In contrast, downregulated terms included fatty acid metabolic processes (GO:0006631, 1.18E-15, group p value Bonferroni corrected), response to hormone (GO:0009725, 7.75E-16, group p value Bonferroni corrected), and nucleobase-containing small molecule metabolism (GO:0055086, 9.42E-14, group p value Bonferroni corrected), consistent with a shift away from catabolic and stress-associated processes. These results suggest that meclizine selectively promotes mitochondrial remodeling and structural recovery while attenuating metabolic stress responses in SLC25A3-deficient hearts.

### Meclizine restores mitochondrial membrane organization in SLC25A3-deficient hearts

As described above, proteomic profiling revealed that meclizine treatment of *Slc25a3^fl/flxMCM^* animals led to upregulation of pathways associated with mitochondrial membrane organization and protein import. Among DEPs in this pathway, we found that meclizine increased expression of five out of the seven subunits of the MICOS complex (MIC13, MIC19, MIC25, MIC26 and MIC27), a critical regulator of cristae junction formation and inner mitochondrial membrane structure (Naha *et al*, 2024; Stephan *et al*, 2020), as well as subunits of the TIM and TOM complexes involved in mitochondrial protein import (Kutik *et al*, 2007) (Figure 4A, B, Supplemental Table 5). In addition, we noted increased expression of ATP5I, a subunit of the FO ATP synthase (Walker *et al*, 1991), that is essential for ATP synthase dimer stabilization and maintaining the high curvature of cristae membranes (Blum *et al*, 2019; Hahn *et al*, 2016) (Figure 4A, Supplemental Table 5), suggesting that meclizine induces the activation of mitochondrial membrane remodeling pathways in SLC25A3-deficient hearts.

**Figure 4.**
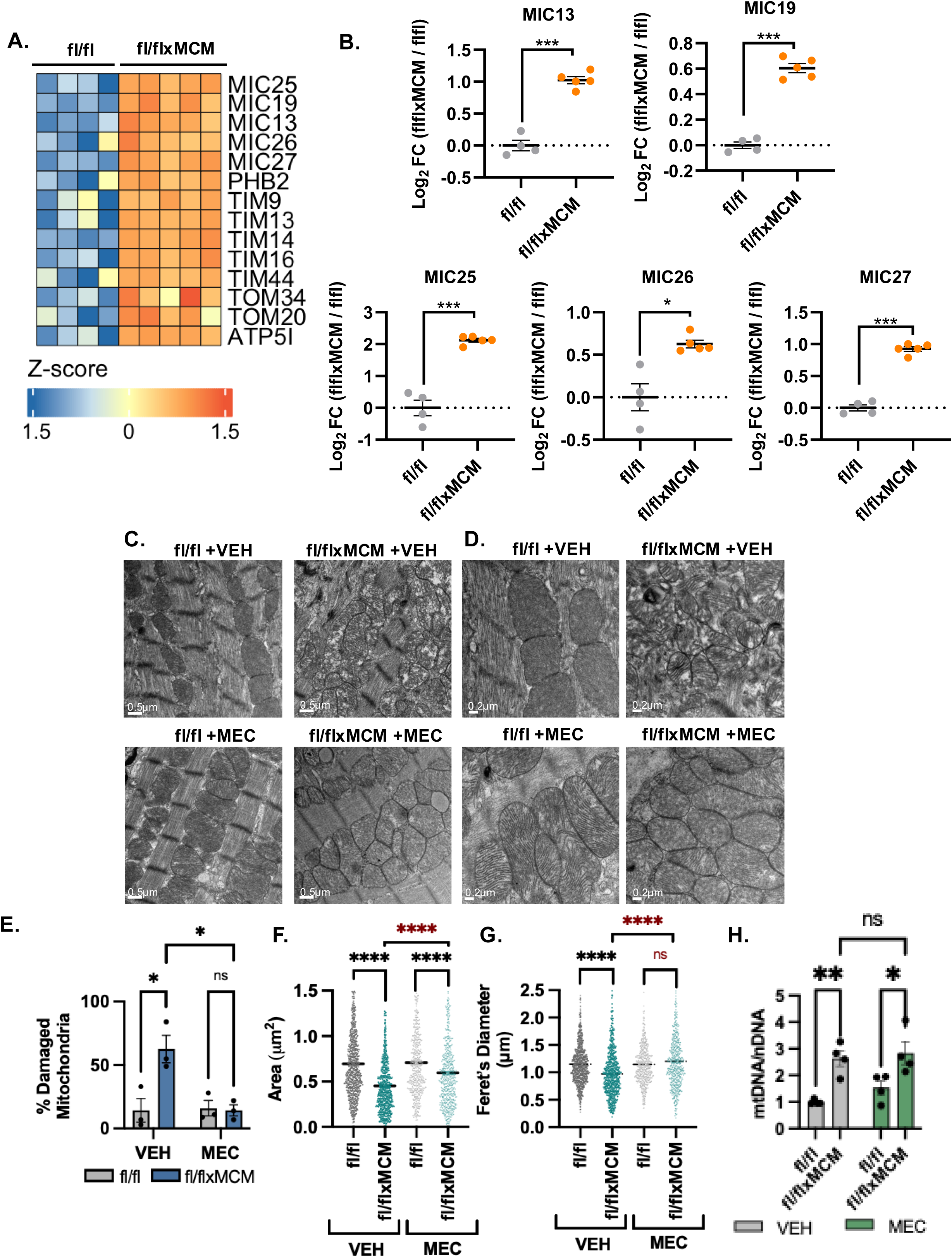
Meclizine ameliorates cristae disorganization and mitochondrial ultrastructural defects in SLC25A3-deficient hearts. (A) TMT proteome levels of MICOS subunits and other mitochondrial membrane organization proteins. All 14 proteins changed more than 1.5-fold in meclizine treated *Slc25a3^fl/flxMCM^*hearts compared to *Slc25a3^fl/fl^* with a *p*-adj < 0.05. Proteins were visualized in a z score normalized heatmap. (B) MS quantification of MIC13, MIC19, MIC25, MIC26, and MIC27 levels in meclizine-treated *Slc25a3^fl/flxMCM^* vs. *Slc25a3^fl/fl^* hearts (Supplemental table 5). (C) Representative transmission electron microscopy (TEM) images from left ventricular tissue from the indicated groups (n=3/group) at magnification X3000. (D) Representative TEM images from left ventricular tissue from the indicated groups (n=3/group) at magnification X5000. (E) Quantitative assessment of damaged mitochondria based on cristae density in the indicated groups (>50 mitochondria were assayed from each replicate n=3/group). (F) Image J quantitative analysis from TEM of mitochondrial area and (G) feret’s diameter from the indicated groups (>300 mitochondria were assayed in each replicate n=3/group). (H) Relative mitochondrial DNA (mtDNA) to nuclear DNA (nDNA) content quantified by qPCR of mitochondrial and nuclear genes (n=4/group). Data are presented as mean ± SEM. For E, F, G and H, two-way ANOVA was used to determine statistical significance \**p* < 0.05, \*\**p* < 0.01, \*\*\**p* < 0.001, \*\*\*\**p* < 0.0001.

To determine if meclizine-associated upregulation mitochondrial membrane organization pathways in SLC25A3-deficient hearts was associated with structural remodeling of the mitochondrial network, we examined cardiac ultrastructure by transmission electron microscopy (TEM). Consistent with our previous findings in these mice (Kwong *et al*., 2014), vehicle-treated *Slc25a3^fl/flxMCM^*hearts displayed hallmark features of mitochondrial cardiomyopathy, including mitochondrial hyperproliferation, fragmentation, cristae disorganization, and disrupted myofibrillar alignment (Figure 4C,D, Supplemental Figure 1, Supplemental Figure 2). Meclizine treatment reversed these structural abnormalities, restoring cristae density and organelle architecture. Quantification of mitochondrial damage showed a reduction from ∼60% damaged mitochondria in vehicle-treated *Slc25a3^fl/flxMCM^* hearts to ∼10% in meclizine-treated hearts (Figure 4E). Morphometric analysis of mitochondria further demonstrated an increase in mitochondrial area and Feret’s diameter in meclizine-versus vehicle-treated *Slc25a3^fl/flxMCM^* hearts, suggesting that meclizine reduces mitochondrial fragmentation and normalizes mitochondrial morphology (Figure 4F, G). In addition to these mitochondrial changes, TEM also revealed that myofibrils in meclizine-treated *Slc25a3^fl/flxMCM^* hearts appeared more organized and better aligned (Supplemental Figure 1), suggesting an overall improvement in cardiac muscle organization.

In addition to assessing mitochondrial ultrastructure, we also evaluated the effects of meclizine on SLC25A3 deletion-associated mitochondrial hyperproliferation. To evaluate this directly, we quantified the mtDNA to nDNA ratio as a measure of mitochondrial content. Intriguingly, while our TEM imaging suggested that meclizine normalized the mitochondrial expansion and improved myofibril alignment in *Slc25a3^fl/flxMCM^* deleted hearts (Figure 4C,D, Supplemental Figure 1, Supplemental Figure 2), mtDNA to nDNA quantification revealed no changes in vehicle versus meclizine treated groups (Figure 4H). Together, these findings suggest that meclizine improves mitochondrial membrane organization, and restores ultrastructural integrity without altering overall mitochondrial abundance in SLC25A3-deficient hearts.

### Meclizine suppresses compensatory glycolysis in SLC25A3 deficient hearts

Given the striking changes to mitochondrial ultrastructure, we next evaluated the impact of meclizine on cardiac energetics. We first assessed the effects of meclizine on mitochondrial function in SLC25A3-deleted and control hearts. As anticipated, mitochondrial respiration and ATP synthesis capacity were decreased in cardiac mitochondria isolated from vehicle-treated *Slc25a3^fl/flxMCM^*versus vehicle-treated *Slc25a3^fl/fl^* control mice, and remained unchanged with meclizine treatment (Figures 5B,C). This was expected since SLC25A3 is essential for mitochondrial ATP synthesis; given the structural defect in the mitochondrial ATP synthesis machinery due to SLC25A3 deletion, meclizine would not be predicted to restore respiration or ATP production.

**Figure 5.**
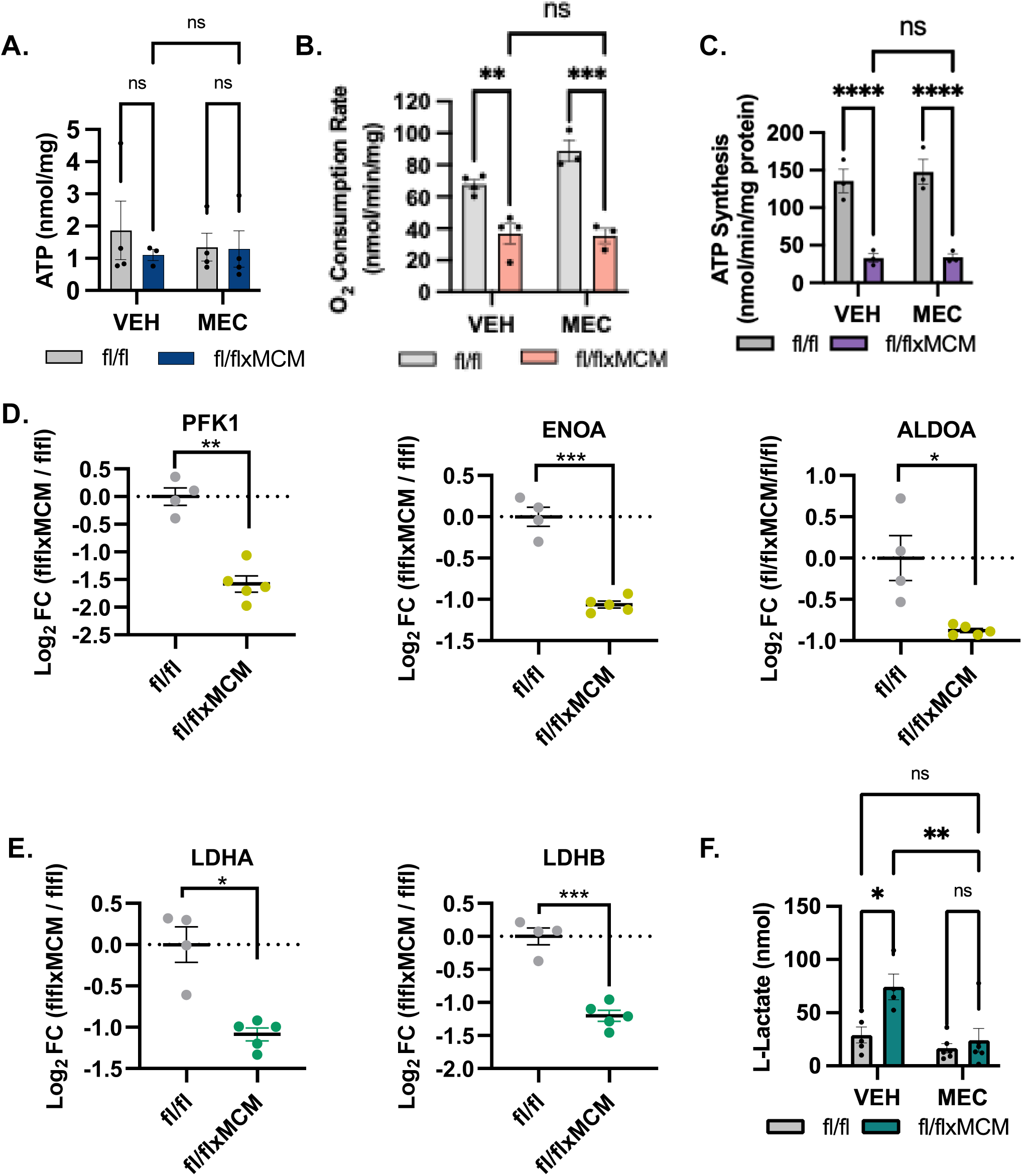
Meclizine suppresses glycolytic compensation despite preserved cardiac ATP levels in SLC25A3-deficient hearts. (A) Total cardiac ATP content (n=4/group). (B) Mitochondrial oxygen consumption rates measured in isolated cardiac mitochondria of the indicated groups (n=3-4/group). (C) Mitochondrial ATP synthesis rate in isolated cardiac mitochondria from the indicated groups (n=3/group). (D) MS quantification of glycolytic enzymes PFK1, ENOA, and ALDOA levels in meclizine-treated *Slc25a3^fl/flxMCM^* vs. *Slc25a3^fl/fl^* hearts (n=5 *Slc25a3^fl/flxMCM^*, n=4 *Slc25a3^fl/fl^*). (E) MS quantification of LDHA and LDHB levels in meclizine-treated *Slc25a3^fl/flxMCM^* vs. *Slc25a3^fl/fl^* hearts (n=5 *Slc25a3^fl/flxMCM^*, n=4 *Slc25a3^fl/fl^*). (F) Measured L-lactate levels in heart tissue from the indicated groups (n=5/group). Data are presented as mean ± SEM. For A, B, C, and F, two-way ANOVA was used to determine statistical significance \**p* < 0.05, \*\**p* < 0.01, \*\*\**p* < 0.001, \*\*\*\**p* < 0.0001. For TMT proteomics quantification of ontologically selected proteins (D and E), differential expression is significant with p-adj < 0.05 and a fold of change of 1.5 (t test followed by Benjamini–Hochberg FDR correction).

Consistent with our previous findings, despite impaired mitochondrial energy production (Figures 5B,C) total cardiac ATP levels were preserved in vehicle-treated *Slc25a3^fl/flxMCM^* hearts versus *Slc25a3^fl/fl^* controls (Figure 5A). However, meclizine treatment did not further increase ATP content of the *Slc25a3^fl/flxMCM^* hearts (Figure 5A), suggesting its effects are not mediated by increasing overall cardiac energy production.

Based on prior studies suggesting that meclizine enhances glycolysis (Gohil *et al*., 2010; Gohil *et al*., 2013; Hong *et al*, 2016; Zhuo *et al*, 2016), we hypothesized that drug treatment would support cardiac energetics under conditions of mitochondrial dysfunction by promoting glycolytic ATP production. Surprisingly, glycolysis was not enhanced. Proteomic analysis revealed that meclizine significantly downregulated the expression of multiple glycolytic enzymes, including aldolase A (ALDOA), enolase 1 (ENO1), and phosphofructokinase-1 (PFK1) in *Slc25a3^fl/flxMCM^*hearts (Figure 5D), indicating suppression of the glycolytic pathway. Moreover, the expression of both LDHA and LDHB, subunits of the lactate dehydrogenase complex responsible for the bidirectional interconversion of pyruvate and lactate, were significantly downregulated with meclizine treatment (Figure 5E), and tissue lactate levels, which were markedly elevated in vehicle-treated *Slc25a3^fl/flxMCM^*hearts, were significantly reduced with meclizine treatment (Figure 5F), further supporting meclizine suppression of SLC25A3 deletion-induced glycolytic compensation.

Taken together, the meclizine-dependent suppression of glycolysis with the maintenance of total cardiac ATP levels in the SLC25A3-deleted mice suggested that meclizine-promoted preservation of cardiac energy balance might occur through a mechanism independent of enhanced glycolytic output.

### Meclizine restores NAD^+^/NADH balance in the SLC25A3-deficient heart by promoting NAD^+^ redox-supportive pathways

Our unexpected results prompted us to further investigate alternative regulatory pathways activated by meclizine that support cardiac energetics in the setting of mitochondrial energy dysfunction. Analysis of our proteomic data revealed that meclizine treatment caused a significant downregulation of pyruvate dehydrogenase kinases PDK2 and PDK4 in *Slc25a3^fl/flxMCM^* hearts as compared to *Slc25a3^fl/fl^* controls (Figure 6A). As PDK2 and PDK4 inhibit pyruvate dehydrogenase to limit the entry of glycolytic carbon into the TCA cycle (Patel & Korotchkina, 2006), downregulation of these kinases would be expected to favor increased pyruvate oxidation and support mitochondrial carbon flux, even though mitochondrial energy output remains unchanged (Figure 5B,C).

**Figure 6.**
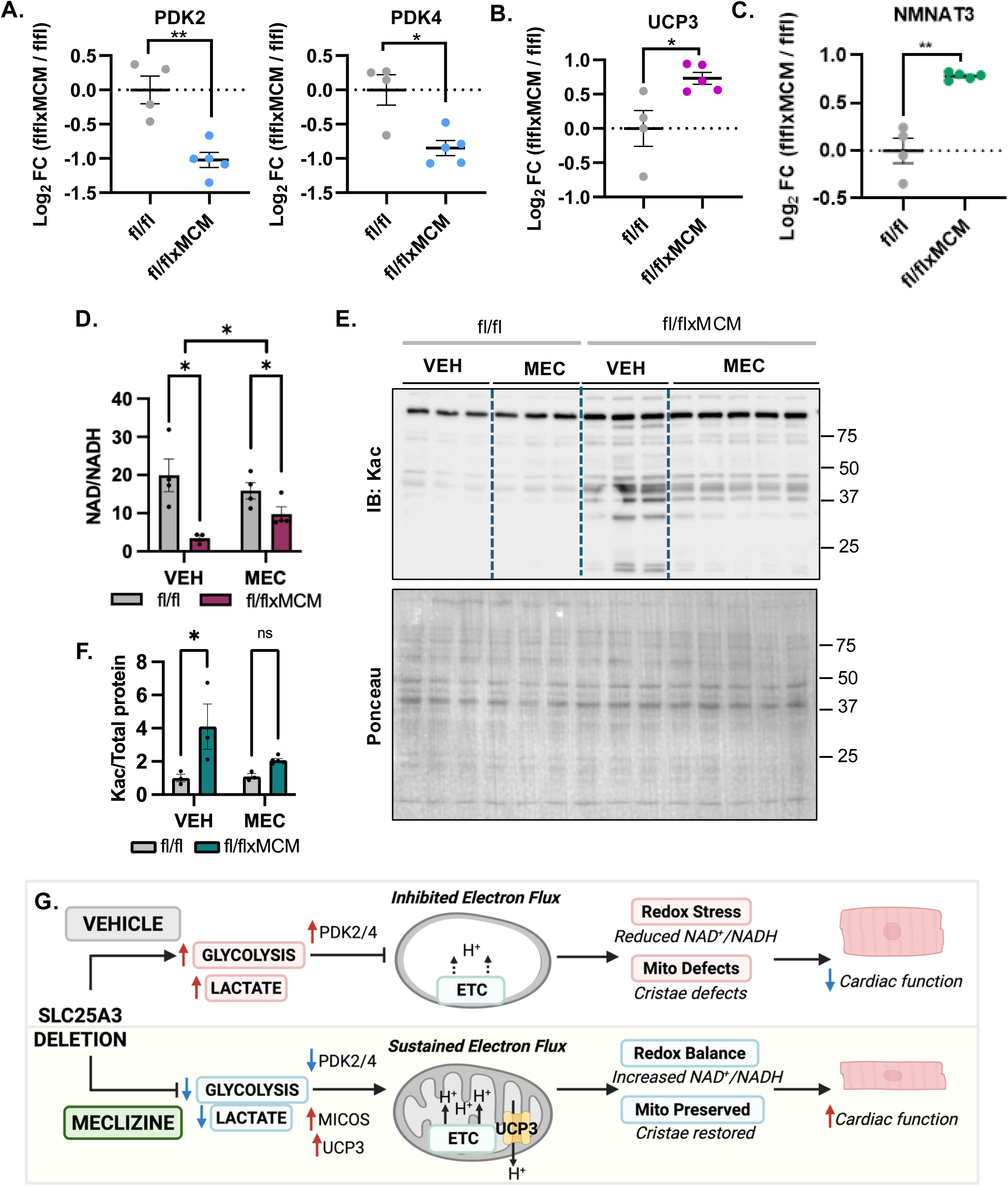
Meclizine restores cardiac redox balance via mitochondrial electron flux and NAD^+^ regeneration. (A) MS quantification of PDK2 and PDK4 in meclizine treated *Slc25a3^fl/flxMCM^* vs *Slc25a3^fl/fl^* hearts (n=5 *Slc25a3^fl/flxMCM^*, n=4 *Slc25a3^fl/fl^*). (B) MS quantification of UCP3 levels in meclizine treated *Slc25a3^fl/f^ ^xMCM^* vs *Slc25a3^fl/fl^* hearts (n=5 *Slc25a3^fl/flxMCM^*, n=4 *Slc25a3^fl/fl^*). (C) MS quantification of NMNAT3 levels in meclizine treated *Slc25a3^fl/f^ ^xMCM^* vs *Slc25a3^fl/fl^* hearts (n=5 *Slc25a3^fl/flxMCM^*, n=4 *Slc25a3^fl/fl^*). (D) Tissue NAD⁺/NADH ratio from the indicated groups (n=4/group). (E) Western blot analyses of acetylated lysines (Kac) in total heart protein lysates prepared from vehicle and meclizine treated *Slc25a3^fl/flxMCM^* and *Slc25a3^fl/fl^* hearts; ponceau stain was used as a loading control. (F) Densitometric quantification of acetyl-lysine (Kac) signal, normalized to total protein loading. Data are presented as mean ± SEM. For TMT proteomics quantification of ontologically selected proteins (A, B, and C), differential expression is significant with p-adj < 0.05 and a fold of change of 1.5 (t-test followed by Benjamini–Hochberg FDR correction). For D and F, two-way ANOVA was used to determine statistical significance \**p* < 0.05, \*\**p* < 0.01, \*\*\**p* < 0.001, \*\*\*\**p* < 0.0001 (G) Proposed model for the effects of meclizine in SLC25A3-deficient hearts. In the absence of SLC25A3, glycolysis and lactate production are elevated, accompanied by upregulation of PDK2/4, which inhibits mitochondrial electron flux. The resulting mitochondria display hyperproliferation, cristae disruption, and redox stress, ultimately leading to cardiomyocyte hypertrophy and impaired cardiac function. In contrast, meclizine treatment lowers glycolysis and lactate production, reduces PDK2/4 expression to relieve inhibition of electron flux, and increases UCP3 expression to dissipate proton buildup, maintaining redox balance. Upregulation of MICOS subunits preserves mitochondrial structure and restores cristae integrity, while reduced TFAM expression indicates decreased mitochondrial hyperproliferation. Together, these effects preserve cardiac function and limit hypertrophy. Diagram created with BioRender.

We postulated that instead of enhancing energy production, meclizine may promote electron flux through the respiratory chain to support redox balance. Interestingly, we observed that meclizine significantly upregulated uncoupling protein 3 (UCP3) in *Slc25a3^fl/flxMCM^* hearts (Figure 6B), which may promote uncoupled electron transport across the inner mitochondrial membrane as a mechanism to sustain electron transport independent of mitochondrial ATP synthesis and could help regenerate NAD^+^ from NADH oxidation to support NAD^+^ redox balance. Consistent with this mechanism, we found that the NAD^+^/NADH ratio was significantly reduced in vehicle treated *Slc25a3^fl/flxMCM^*versus *Slc25a3^fl/fl^* control hearts, but this ratio was significantly increased with meclizine treatment (Figure 6D). In line with these changes, we found that nicotinamide mononucleotide adenylyltrasferase3 (NMNAT3) was significantly upregulated with meclizine (Figure 6C). Because NMNAT3 is responsible for the conversion of nicotinamide mononucleotide to NAD⁺ within mitochondria (Berger *et al*, 2005), upregulation of NMNAT3 may provide an additional route for maintaining mitochondrial NAD⁺ pools under stress conditions. In further support of meclizine-induced improvement of cardiac NAD^+^ redox status, meclizine-treated *Slc25a3^fl/flxMCM^* hearts also exhibited significantly reduced protein hyperacetylation compared to vehicle-treated *Slc25a3^fl/flxMCM^* hearts (Figure 6E,F), a change consistent with increased NAD⁺ availability to support the activity of NAD⁺-dependent protein deacetylases (Lee et al., 2016). Together, these findings suggest that meclizine promotes cardiac energy homeostasis not by increasing ATP production, but by activating a metabolic circuit that supports NAD⁺ regeneration and NAD/NADH balance (Figure 6G). This NAD^+^ redox-supportive adaptation likely contributes to the preservation of cardiac function in the face of intrinsic mitochondrial ATP synthesis defects caused by SLC25A3 deficiency.

## DISCUSSION

Mitochondrial cardiomyopathies remain a significant clinical challenge due to the lack of targeted therapies that address the underlying mitochondrial dysfunction (Meyers *et al*., 2013). To address the need for interventions that can support cardiac energy supply despite mitochondrial energy dysfunction, we defined the cardiac impact of the FDA-approved compound meclizine, which has been reported to promote a metabolic shift toward glycolysis (Gohil *et al*., 2010; Gohil *et al*., 2013; Hong *et al*., 2016; Zhuo *et al*., 2016). We demonstrate that meclizine confers cardioprotection in a mouse model of mitochondrial cardiomyopathy caused by cardiomyocyte-specific deletion of SLC25A3, a mitochondrial inner membrane transporter essential for inorganic phosphate import and, thus, mitochondrial ATP synthesis. Chronic systemic meclizine treatment attenuated cardiac hypertrophy, improved systolic function, and restored mitochondrial ultrastructure. As expected, meclizine did not correct the mitochondrial respiration or ATP synthesis defects caused by SLC25A3 loss. Surprisingly, however, meclizine also failed to enhance glycolysis, suggesting an alternative mechanism of meclizine action.

Indeed, a key unexpected finding of this study is that meclizine suppresses, rather than enhances, glycolytic compensation in SLC25A3-deficient hearts. In previous work, we reported that SLC25A3 deletion impairs mitochondrial ATP synthesis but does not affect total cardiac ATP levels, an effect we had attributed to increased glucose uptake and upregulation of glycolytic enzymes (Kwong *et al*., 2014). Those observations led us to postulate that therapeutic strategies that promote a compensatory switch to glycolytic metabolism would be protective in the context of intrinsic defects in the mitochondrial energy production machinery. However, contrary to prior reports (Gohil *et al*., 2010; Gohil *et al*., 2013; Hong *et al*., 2016) and our hypothesis, meclizine treatment in the *Slc25a3^fl/flxMCM^* mice downregulated the expression of key glycolytic enzymes (ALDOA, ENO1, PFK1). Additionally, the lactate dehydrogenase subunits LHDA and LDHB, were downregulated. Both LDHA and LDHB can catalyze the bidirectional conversion of pyruvate to lactate with LDHA showing a higher affinity for the conversion of pyruvate to lactate and LDHB possessing a higher affinity for the reverse reaction (Urbanska & Orzechowski, 2019). The concurrent downregulation of both subunits together with reduced tissue lactate levels, suggests a broader dysregulation of lactate metabolism. Importantly, while decreased LDHA aligns with suppressed glycolytic flux, reduced LDHB, which is highly expressed in the heart and facilitates lactate oxidation (Tian *et al*, 2020), may indicate impaired utilization of lactate as a metabolic substrate. Thus, meclizine appears to disrupt not only glycolysis but also lactate homeostasis in the SLC25A3-deficient heart.

Instead, our findings support a model in which meclizine protects the SLC25A3-deleted heart in part by preserving mitochondrial architecture. In line with our previous observations, we found that vehicle-treated *Slc25a3^fl/flxMCM^*hearts displayed aberrant mitochondria characterized by disrupted cristae, fragmentation, and pathological hyperproliferation. Quantitative proteomics comparing the effects of meclizine versus vehicle treatment of *Slc25a3^fl/flxMCM^*animals revealed that meclizine treatment caused a robust upregulation of MICOS complex subunits, including MIC13, MIC19, MIC25, MIC26, and MIC27, along with components of the TIM and TOM import machinery, pathways that are important for maintaining cristae architecture and mitochondrial integrity (Harner *et al*, 2011; Hoppins *et al*, 2011; Huynen *et al*, 2016; Korner *et al*, 2012; Stephan *et al*., 2020; von der Malsburg *et al*, 2011). Indeed, these proteomic changes were supported by TEM, which showed that meclizine restored normal cristae morphology, reduced mitochondrial fragmentation, normalized mitochondrial size, and improved myofibrillar alignment in SLC25A3-deficient hearts. Taken together, these data highlight mitochondrial membrane organization as a key therapeutic target of meclizine, linking improved mitochondrial ultrastructure with improved cardiomyocyte organization that may be beneficial even in the absence of restored mitochondrial ATP synthesis.

Notably, meclizine treatment also promoted NAD^+^ redox balance and mitochondrial metabolic flux despite the suppression of the compensatory glycolysis in the SLC25A3-deleted hearts. Our proteomic analysis revealed that meclizine downregulated the pyruvate dehydrogenase kinases PDK2 and PDK4, which inhibit the pyruvate dehydrogenase complex to restrict carbon entry into the TCA cycle. This suggests that meclizine facilitates increased pyruvate oxidation and carbon flow through mitochondrial pathways, even though mitochondrial ATP synthesis remains impaired. Indeed, mitochondrial respiration and ATP production were not restored by meclizine, indicating that increased substrate flow does not enhance mitochondrial energy output under conditions of SLC25A3 loss.

Instead, our data suggest that meclizine promotes NAD^+^ regeneration to support NAD^+^ redox homeostasis. Meclizine-treated *Slc25a3^fl/flxMCM^*hearts showed increased expression of UCP3. While originally described as an uncoupling protein that mediates proton leak (Boss *et al*, 1997; Krauss *et al*, 2005), UCP3 has been suggested to have broader functions, including facilitating fatty acid anion export, mitigating lipid peroxidation as well as regulating ROS production (Pohl *et al*, 2019). In the setting of SLC25A3 deletion, meclizine-induced increases in UCP3 may offer a pathway of uncoupled electron transport to enable oxidation of NADH to NAD^+^, helping maintain a favorable NAD^+^/NADH ratio. Supporting this model, we observed a significant increase in the NAD^+^/NADH ratio in meclizine-versus vehicle-treated *Slc25a3^fl/flxMCM^* hearts.

In parallel, we also found that meclizine upregulated NMNAT3 in *Slc25a3^fl/flxMCM^*hearts. NMNAT3 is a key mitochondrial enzyme in the NAD⁺ salvage pathway that catalyzes the conversion of nicotinamide mononucleotide (NMN) to NAD^+^, which could further contribute to the observed meclizine-associated increase in NAD^+^/NADH ratio (Berger *et al*., 2005). These findings suggest that rather than restoring energy production, meclizine enables the heart to maintain redox stability through coordinated regulation of mitochondrial carbon flux and NAD^+^ regeneration, which aligns with studies demonstrating that increasing cardiac NAD^+^/NADH ratio via the NAD^+^ salvage pathway by supplementation with nicotinamide riboside or NMN, precursors of NAD^+^ synthesis, improves cardiac function and stress tolerance in models of heart failure (Diguet *et al*, 2018; Lee *et al*, 2019; Lee *et al*, 2016). Our work extends these observations by identifying meclizine as a clinically approved compound that converges on similar NAD^+^ regulatory pathways while also ameliorating mitochondrial structure. However, whether meclizine-mediated improvements in mitochondrial ultrastructure and NAD^+^ redox balance represent independent protective mechanisms or are mechanistically linked remains to be determined. It is possible that stabilization of cristae architecture and restoration of MICOS complex expression enhance respiratory chain organization and facilitate electron transport in SLC25A3 deleted mitochondria, thereby improving NAD^+^ redox status. Alternatively, improved NAD⁺ availability and NAD^+^/NADH ratio may secondarily promote mitochondrial membrane integrity and suppress maladaptive remodeling. Future studies are needed to delineate the temporal and causal relationships among these pathways and to determine whether they represent components of a single regulatory axis or distinct elements of a multifaceted protective response to mitochondrial energy dysfunction.

In summary, meclizine engages a coordinated protective response involving improved mitochondrial structure and redox homeostasis in the context of SLC25A3-associated mitochondrial cardiomyopathy, though the extent to which these pathways might be mechanistically linked remains to be defined. Meclizine had no significant effects in the control heart; yet, in the context of the SLC25A3-deleted heart, meclizine appears to coordinate an integrated cardioprotective program by preserving mitochondrial membrane architecture, supporting redox balance through NAD^+^ regeneration, and reducing cardiac protein acetylation to allow the heart to maintain contractile function despite defects in the mitochondrial energy production machinery. These findings are consistent with the demonstrated protective effects of enhancing NAD^+^ in the heart and highlight the therapeutic potential of targeting both mitochondrial quality and redox homeostasis in mitochondrial disease. While further work is needed to understand how meclizine engages these pathways, the optimal therapeutic window for meclizine cardioprotection, if meclizine’s protective effects can extend to mitochondrial cardiomyopathy due to other types of primary mitochondrial defects or even common cardiac diseases with mitochondrial dysfunction like heart failure, our results identify meclizine as a promising candidate for promoting mitochondrial health in in the heart.

## METHODS

### Animals

Conditional knockout mice with loxP-flanked *Slc25a3* alleles (*Slc25a3^fl/fl^*) and transgenic mice expressing a tamoxifen-inducible Cre recombinase under the control of the cardiomyocyte-specific α-myosin heavy chain promoter (αMHC-MerCreMer, or MCM) were previously described (Kwong *et al*., 2014). To generate cardiac-specific *Slc25a3* knockout mice, *Slc25a3^fl/fl^* animals were bred with MCM mice as described (*Slc25a3^fl/fxMCM^*) (Kwong *et al*., 2014; Peoples *et al*., 2021). The MCM transgene was consistently maintained in the hemizygous state. At 8 weeks of age, *Slc25a3^fl/fxMCM^* mice or *Slc25a3^fl/fl^*controls received daily intraperitoneal injections of tamoxifen (25 mg/kg) for five consecutive days to induce *Slc25a3* deletion. Concurrently with tamoxifen administration, mice were randomized into either vehicle groups [10% Kolliphor (Sigma-Aldrich)] or meclizine groups [100 mg/kg/day meclizine dihydrochloride (Santa Cruz Biotechnology) by oral gavage] that received treatment for 12 weeks. Both male and female mice were included in the study. Experimental analyses were conducted at 12 weeks following tamoxifen treatment. For tissue collection, mice were anesthetized with isoflurane and euthanized via cervical dislocation. All animal procedures were reviewed and approved by the Institutional Animal Care and Use Committee (IACUC) at Emory University.

### Echocardiography

Mice were anesthetized using 1.5% isoflurane, with body temperature maintained at 37°C during the procedure. Cardiac imaging was performed using a Vevo 3100 Imaging System (VisualSonics) equipped with an MS-250 transducer. M-mode images were acquired from the parasternal short axis view. Parameters measured using VevoLab software included left ventricular internal diameter in diastole (LVID,d) and systole (LVID,s), left ventricular volume in diastole (V,d) and systole (V,s), stroke volume (SV), fractional shortening (FS), and ejection fraction (EF).

### Immunoblotting

For Western blot analyses, total protein extracts were prepared from hearts homogenized and solubilized in radioimmunoprecipitation assay buffer supplemented with 2.4 mg/mL nicotinamide, 500 nM trichostatin A and a combined protease and phosphatase inhibitor cocktail (Thermo Fisher Scientific). For electrophoresis, proteins were reduced and denatured in Laemmli buffer, resolved on 10% SDS-PAGE gels, transferred to PVDF membranes, immunodetected with antibodies, and imaged using a ChemiDoc system (BioRad). Primary antibodies used in the study were anti-acetylated lysine (PTM Biolabs PTM-105, 1:1,000). Secondary antibodies were alkaline phosphatase-linked goat antirabbit IgG (Cell Signaling Technologies #7054, 1:5,000), alkaline phosphatase-linked goat anti-mouse IgG (Cell Signaling Technologies #7056, 1:5,000), and horseradish peroxidase-linked goat anti-mouse IgG (Cell Signaling Technologies #7076, 1:5,000).

### TMT-Based Quantitative Proteomics

Mouse heart tissue was collected, snap frozen in liquid nitrogen, and stored at −80°C until further processing. 30 mg of tissue per sample was homogenized in modified RIPA buffer (50 mM Tris-HCl, pH 8.0, 150 mM NaCl, 2.0% SDS, 0.1% Triton X-100) supplemented with Roche Complete protease inhibitors (Millipore-Sigma). Samples were sonicated using a probe sonicator (35–40 % amplitude, 1 s on/off pulses for 20 s total), clarified by centrifugation at 13,200 rpm for 15 min at 4°C, and protein concentration was determined using the Qubit Protein Assay Kit (Thermo Fisher Scientific). Proteins were precipitated with trichloroacetic acid (TCA) overnight at −20°C. Pellets were washed with cold acetone, air-dried, and resolubilized in 8 M urea, 50 mM Tris-HCl, pH 8.0 containing protease inhibitors. A total of 50 µg of protein per sample was subjected to tryptic digestion. Proteins were reduced in 12 mM DTT for 1 h at room temperature, followed by alkylation in 15 mM iodoacetamide for 1 h in the dark. Urea was diluted to 1 M with 50 mM TEAB, and sequencing-grade modified trypsin (Promega) was added at an enzyme-to-substrate ratio of 1:20. Samples were incubated overnight at 37°C. Digestion was stopped by acidifying to 0.3% trifluoroacetic acid (TFA). Peptides were desalted by solid-phase extraction (SPE) using Waters µElution HLB plates (Waters, #186001828BA). Columns were conditioned with 4 × 500 µL of 70% acetonitrile (ACN), equilibrated with 4 × 500 µL of 0.3% TFA, and the acidified digests were loaded. Columns were washed with 3 × 500 µL of 0.3% TFA and eluted in two steps: 200 µL followed by 400 µL of 60% ACN in 0.3% TFA. Eluates were frozen at −80°C and lyophilized overnight. Dried peptides were resuspended in 140 mM HEPES, pH 8.0, 30% ACN for TMT labeling. Each TMTpro reagent (Thermo Fisher Scientific) was reconstituted in 40 µL anhydrous ACN, incubated for 15 min at room temperature, and 30 µL was added to the corresponding peptide sample. Labeling reactions were incubated at 25°C for 1.5 h in an Eppendorf Thermomixer at 300 rpm. Labeling was quenched by adding 8 µL of 5% hydroxylamine and incubating for 15 min. A ratio check was performed by pooling 1% of each sample into a single 18-plex, followed by desalting and LC-MS analysis to confirm relative channel intensities. Based on this, samples were normalized, pooled accordingly, lyophilized, and subjected to another round of SPE cleanup using Waters HLB cartridges (WAT106202). Fifty percent of the pooled sample was fractionated by high-pH reverse-phase liquid chromatography. Peptides were resuspended in 150 µL of Buffer A (10 mM ammonium hydroxide, pH 10.5 in water) and injected onto an XBridge BEH C18 column (2.1 × 150 mm, 3.5 µm; Waters, #186003023) using an Agilent 1100 HPLC system operating at 0.3 mL/min with UV detection at 214 nm. The following gradient was applied: 0–1 min, 0.5% B; 1–36 min, linear to 25% B; 36–44 min, linear to 45% B; 44–47 min, ramp to 90% B; 47–49 min, hold at 90% B; 49–50 min, re-equilibration at 0.5% B (Buffer B: 10 mM ammonium hydroxide, pH 10.5 in ACN). Ninety-six 150 µL fractions were collected at 30 s intervals and concatenated into 24 pools using a checkerboard pattern. Fractions were dried by vacuum centrifugation and stored at −80°C. Peptides from each pooled fraction were analyzed by nanoLC-MS³ using a Waters NanoAcquity UPLC coupled to a ThermoFisher Fusion Lumos mass spectrometer. Samples were loaded onto a trapping column and separated on a 75 µm i.d. analytical column packed in-house with Luna C18 resin (Phenomenex). Elution was performed over a 1.5 h gradient at 350 nL/min. The MS³ method included full MS scans in the Orbitrap at 120,000 resolution, followed by MS² CID scans in the ion trap with normalized collision energy (NCE) of 32%, and MS³ HCD scans in the Orbitrap at 50,000 resolution from m/z 100–500. Synchronous precursor selection (SPS) was used to isolate up to 10 MS² fragment ions for MS³ quantification. A 2s top-speed cycle time was applied.

### TEM and mitochondrial ultrastructural analysis

Small blocks of mouse heart tissue (1 mm³) were fixed in 2.5% glutaraldehyde in 0.1 M sodium cacodylate buffer (pH 7.4), post-fixed with 1% osmium tetroxide, dehydrated, and embedded in epoxy resin. Ultrathin sections (80-90 nm) were cut and stained with uranyl acetate and lead citrate and imaged using a JEOL 1400 transmission electron microscope (Tokyo, Japan) equipped with a Gatan US1000 CCD camera (Pleasanton, CA). For mitochondrial morphometric analysis, 100–200 mitochondria were analyzed per sample (n=3/group) using Fiji (ImageJ) software. Mitochondrial area and Feret’s diameter were quantified using threshold-based segmentation and particle analysis tools.

### mtDNA Copy Number Analysis

Frozen heart tissue was used to extract total DNA using the Qiagen DNeasy Blood & Tissue Kit (#69504). DNA purity and concentration were measured with NanoDrop. Quantification of relative mtDNA copy number was conducted by measuring mtDNA/nDNA ratio using qPCR qPCR was performed using SYBR Green Master Mix (Biorad) on a CFX Maestro software. Primers for mouse mtDNA (forward, CTAGAAACCCCGAAACCAAA; reverse, CCAGCTATCACCAAGCTCGT) and the β-2-microglobulin nuclear DNA (forward, 5’ ATGGGAAGCCGAACATACTG 3’; reverse, 5’ CAGTCTCAGTGGGGGTGAAT 3’) as described previously (Ghazal *et al*, 2021). Relative mtDNA copy number was calculated using the ΔΔCt method.

### Mitochondrial isolation

Cardiac mitochondria were isolated by differential centrifugation as described (Ghazal *et al*., 2021). Briefly, freshly excised hearts were minced in ice-cold MS-EGTA buffer (225 mM mannitol, 75 mM sucrose, 10 mM HEPES, 1 mM EGTA, pH 7.4). Tissues were homogenized using a Teflon-glass Dounce homogenizer (10 passes). Homogenates were centrifuged at 600 × g for 5 min at 4°C to remove debris and nuclei. Supernatants were centrifuged at 10,000 × g for 10 min. Pellets were washed and resuspended in MS-EGTA buffer, then centrifuged again at 10,000 × g for 10 min. Final mitochondrial pellets were resuspended in 50–100 µL of isolation buffer.

### ATP measurements

Total ATP levels in whole heart lysates were measured using a luciferase-based ATP determination kit (Invitrogen) according to the manufacturer’s protocol. For mitochondrial ATP synthesis, a modified protocol was used for mitochondrial ATP synthesis as described previously (Kwong *et al*., 2014). Briefly 50 mg isolated heart mitochondria were incubated with 0.15 mM adenosine pentaphosphate, 1 mM malate, 1 mM glutamate, and 0.1 mM ADP in Buffer A (150 mM KCL, 25 mM Tris-HCl, 2 mM EDTA, 0.1 % bovine serum albumin, 10 mM K-phosphate, 0.1 mM MgCl, pH 7.4). Buffer B (0.5 M Tris acetate pH7.75, 0.8 mM luciferin, and 20 µg/ml luciferase) was added to the mitochondria and luminescence was measured in kinetic mode every 10 s for 4 min using a Synergy Neo Multimode plate reader (BioTek).

### Lactate assay

Cardiac lactate levels were measured using the L-Lactate Assay Kit (Abcam, ab65330) following the manufacturer’s protocol. Briefly, 10–20 mg of frozen heart tissue was minced and homogenized using a glass homogenizer in 300 µL of assay buffer. Samples were deproteinized using perchloric acid (PCA) precipitation followed by neutralization with potassium hydroxide (KOH), and the resulting supernatants were collected for analysis. For quantification, samples were incubated with lactate assay reaction mix for 30 minutes at room temperature, and absorbance was measured at 570 nm using a Synergy Neo Multimode reader (BioTek).

### NADH assay

NAD⁺ and NADH levels were quantified using the NAD⁺/NADH Quantification Colorimetric Kit (Abcam, ab65348). Heart tissue (20 mg) was extracted in 400 µL NAD/NADH extraction buffer according to the kit’s protocol. Extracts were split: 200 µl heated at 60°C for 30 min to decompose NAD⁺ (for NADH-only measurement), and the other 200 µl was used for total NADH measurement. After neutralization, samples were incubated with the cycling enzyme mix for 5 min and then with the NADH developer for 1 h. Absorbance was measured at 450 nm using a Synergy Neo Multimode reader (BioTek).

### Respiration

Mitochondrial oxygen consumption rates were measured using an Oxytherm+ R system (Hansatech) as previously described (Ghazal *et al*., 2021). 100 mg of isolated cardiac mitochondria were incubated in respiration buffer composed of 120 mM KCl, 5 mM MOPS, 0.1 mM EGTA, 5 mM KH_2_PO_4_, 0.2 % fatty acid-free BSA, 10 mM glutamate, and 2 mM malate. ADP-stimulated respiration was initiated by adding 0.5 mM ADP. To assess maximal uncoupled respiration, 5 µM FCCP was added. Non-mitochondrial respiration was measured following the addition of potassium cyanide (KCN). Mitochondrial oxygen consumption rates were calculated by subtracting KCN-insensitive values from basal and FCCP-stimulated measurements and normalized to protein content.

### Bioinformatic analyses and statistical analyses

Raw data were analyzed using MaxQuant v1.6.14.0. MS spectra were recalibrated, and database searches were performed using Andromeda against the SwissProt mouse proteome database. Search parameters included: enzyme specificity set to trypsin (cleavage at K/R, not before P), fixed modification of carbamidomethyl (C), and variable modifications of oxidation (M) and acetylation (protein N-terminus). TMT reporter ion intensities were extracted and corrected for isotopic impurities. Search results were filtered to a 1% FDR at both peptide and protein levels.

For TMT quantification, reporter ion intensities were extracted, isotopically corrected using manufacturer-provided factors, and log₂-transformed. Intensities were normalized by subtracting the median value for each TMT channel, and only proteins with valid intensity values in all samples were retained for downstream analysis (2,803 proteins). Perseus software (version 1.5.5.3) was used to calculate group means and prepare processed data matrices.

Gene Ontology (GO) Biological Process enrichment analysis was performed in Cytoscape (version 3.10.2; https://cytoscape.org) using the ClueGO plugin (version 2.5.10) (Bindea *et al*, 2009). Analyses were conducted with the Mus musculus annotation set using two-sided hypergeometric testing with Bonferroni correction (p ≤ 0.05). GO terms between levels 3 and 8 were considered, with a minimum of 3% of genes per term. GO Fusion and grouping were enabled, and terms were clustered based on a Kappa score threshold of 0.4. Functionally related terms were named according to the most significant term in each group.

All additional graphs and statistical comparisons for non-proteomics datasets were generated in GraphPad Prism (version 8; GraphPad Software, San Diego, CA). Data are presented as mean ± standard error of the mean (SEM). Statistical significance between two groups was assessed using two-tailed unpaired Student’s t-tests, unless otherwise specified. Differences were considered statistically significant at *p* ≤ 0.05.

---

**Supplemental Figure 1.**
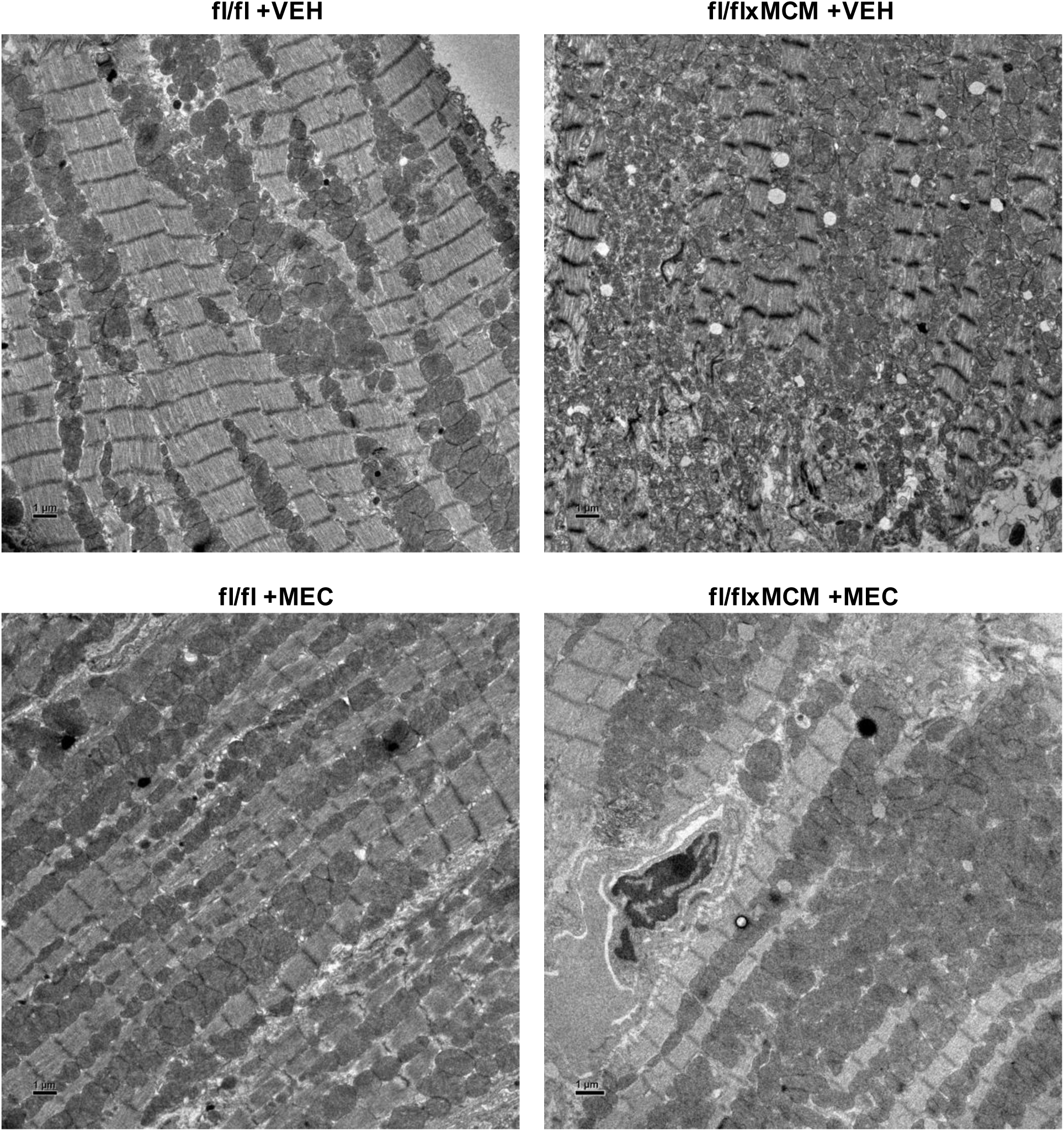
Meclizine reduces mitochondrial hyperproliferation and improves myofibrillar alignment. Transmission electron micrographs of vehicle treated (top) and meclizine treated (bottom) *Slc25a3^fl/fl^* and *Slc25a3^fl/flxMCM^* hearts after 12 weeks of tamoxifen (n=3/group). Vehicle treated *Slc25a3^fl/flxMCM^*hearts show hyperproliferated mitochondria and disrupted myofilament architecture. In contrast, meclizine decreases mitochondrial hyperproliferation and promotes better myofilament alignment.

**Supplemental Figure 2.**
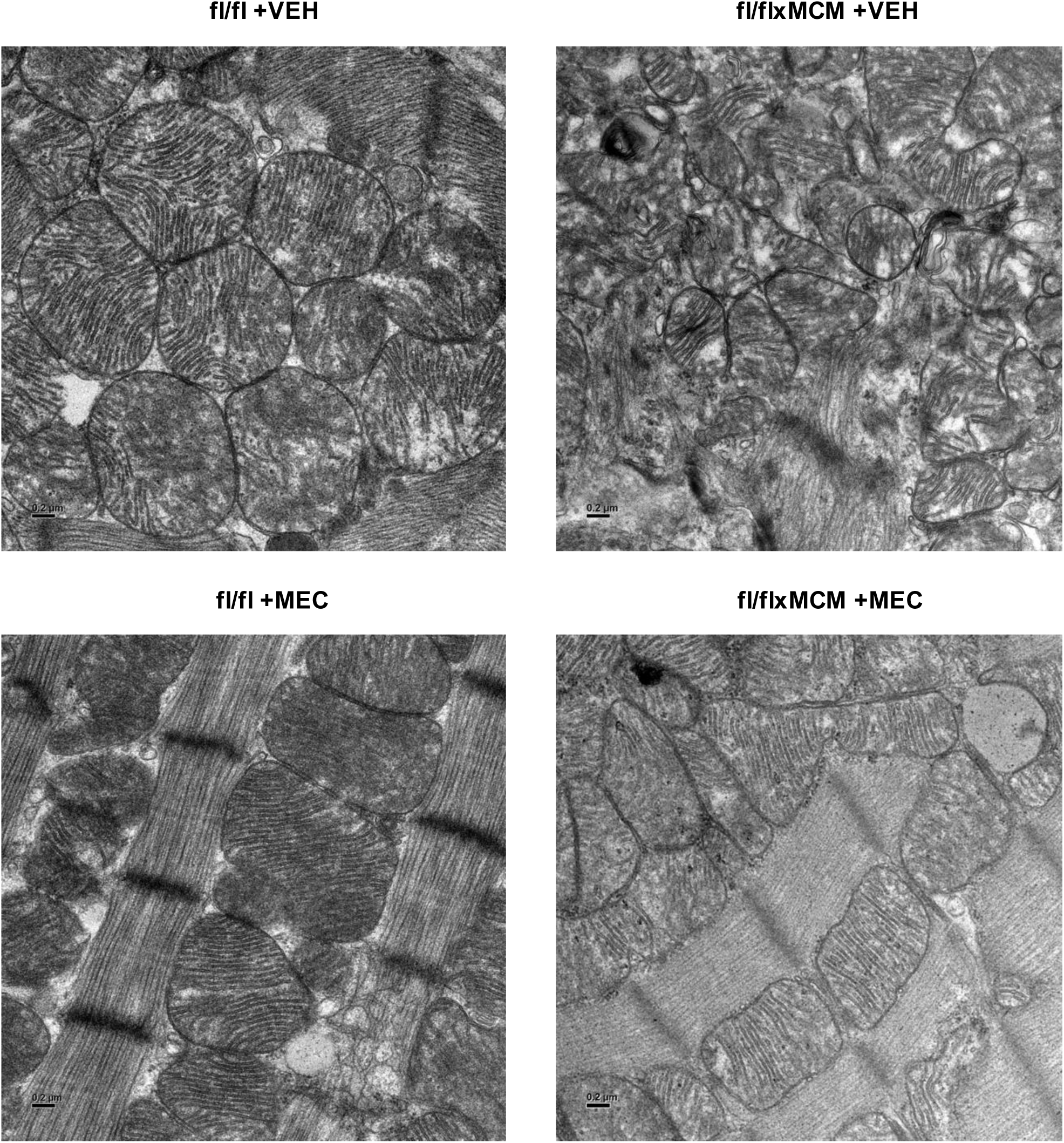
Meclizine improves mitochondrial ultrastructure and restores cristae organization in SLC25A3-deficient hearts. High magnification transmission electron micrographs of vehicle treated (top) and meclizine treated (bottom) *Slc25a3^fl/fl^* and *Slc25a3^fl/flxMCM^* hearts after 12 weeks of tamoxifen (n=3/group). Vehicle treated *Slc25a3^fl/flxMCM^* hearts show fragmented mitochondria with disrupted and sparse cristae. In contrast, meclizine ameliorates mitochondrial fragmentation and restores cristae density and alignment.

**Supplemental Table 1. Complete TMT-based proteomics dataset with normalized abundance values and pairwise comparisons.**

**Supplemental Table 2. Gene Ontology Biological Process enrichment analysis for the vehicle comparison (fl/flxMCM vs fl/fl).**

**Supplemental Table 3. Protein changes in vehicle- and meclizine-treated groups with log₂ fold change and adjusted p-values for Pearson correlation analysis.**

**Supplemental Table 4. Gene Ontology Biological Process enrichment analysis for the meclizine comparison (fl/flxMCM vs fl/fl).**

**Supplemental Table 5. Mitochondrial membrane organization-annotated genes differentially expressed with meclizine treatment.**

## AUTHOR CONTRIBUTIONS

JQK and NG wrote the manuscript. NG, BH, and LJS performed experiments. JQK, NG, and LJS analyzed the data. JQK, NG, and VF designed the study. JQK conducted experimental oversight.

## ACKNOWLEDGEMENTS

The authors thank Austin Park for assisting with ATP synthesis and respiration measurements. JQK was supported by the National Institutes of Health (R01GM144729). NG was supported by an American Heart Association Predoctoral Fellowship (25PRE1372965). VF was supported by the National Institutes of Health (ES034796). Electron microscopy for this study was supported by the Robert P. Apkarian Integrated Electron Microscopy Core, which is subsidized by the Emory University School of Medicine and the Emory College of Arts and Sciences. Additional support for electron microscopy was provided by the Georgia Clinical and Translational Science Alliance of NIH (UL1TR000454). Echocardiography for this study was supported by the Animal Physiology Core, which is subsidized by Emory University and Children’s Healthcare of Atlanta. Additional support was provided by the NIH Office of the Director (S10OD021748).

